# SARS-CoV-2 Spike Protein Induces Hemagglutination: Implications for COVID-19 Morbidities and Therapeutics and for Vaccine Adverse Effects

**DOI:** 10.1101/2022.11.24.517882

**Authors:** Celine Boschi, David E. Scheim, Audrey Bancod, Muriel Millitello, Marion Le Bideau, Philippe Colson, Jacques Fantini, Bernard La Scola

## Abstract

Experimental findings for SARS-CoV-2 related to the glycan biochemistry of coronaviruses indicate that attachments from spike protein to glycoconjugates on the surfaces of red blood cells (RBCs), other blood cells and endothelial cells are key to the infectivity and morbidity of COVID-19. To provide further insight into these glycan attachments and their potential clinical relevance, the classic hemagglutination (HA) assay was applied using spike protein from the Wuhan, Alpha, Delta and Omicron B.1.1.529 lineages of SARS-CoV-2 mixed with human RBCs. The electrostatic potential of the central region of spike protein from these four lineages was studied through molecular modeling simulations. Inhibition of spike protein-induced HA was tested using the macrocyclic lactone ivermectin (IVM), which is indicated to bind strongly to SARS-CoV-2 spike protein glycan sites. The results of these experiments were, first, that spike protein from these four lineages of SARS-CoV-2 induced HA. Omicron induced HA at a significantly lower threshold concentration of spike protein than for the three prior lineages and was much more electropositive on its central spike protein region. IVM blocked HA when added to RBCs prior to spike protein and reversed HA when added afterwards. These results validate and extend prior findings on the role of glycan bindings of viral spike protein in COVID-19. They furthermore suggest therapeutic options using competitive glycan-binding agents such as IVM and may help elucidate rare serious adverse effects (AEs) associated with COVID-19 mRNA vaccines which use spike protein as the generated antigen.

## INTRODUCTION

Key to the infectivity and morbidity of SARS-CoV-2 are glycans that protrude tangentially from 22 N-linked glycosylation sites on each monomer of its spike protein.^1-5^ These N-glycans, several of which are capped with terminal sialic acid (SA) moieties, sweep back and forth across spike protein like windshield wipers, partially shielding it from antibody binding.^1,2,6,7^ For SARS-CoV-2 and several other coronaviruses, these N-glycans serve as appendages for the virus to make its initial attachments to glycoconjugates on the host cell surface.^1,8-13^ These glycoconjugates confer a negative electrostatic potential on the surfaces of host cells such as red blood cells (RBCs),^14,15^ platelets^16^ and endothelial cells,^17^ additionally facilitating attachments by positively charged SARS-CoV-2 spike protein prior to viral fusion to ACE2 for replication.^8,18-20^ For endothelial cells of blood vessel linings, for example, the disparity between 28,000 SA-tipped CD147 receptors and 175 ACE2 receptors per cell^21^ provides a supporting indication of the role of glycans in widespread endothelial damage reported in COVID-19 patients.^22,23^

The RBC has an especially dense surface distribution of SA, 35 million SA molecules per cell, arrayed on its sialoglycoprotein coating mainly as terminal residues of glycophorin A (GPA).^14,15^ SA in its predominant human form, Neu5Ac, is the most common terminal residue of GPA on human RBCs, with the other terminal monosaccharides of GPA matching those on SARS-CoV-2 spike N-glycans.^2,5^ Through attachments to viruses via this GPA surface coating, RBCs and other blood cells can serve a host defense role,^14,24,25^ however these RBC clumps can be vascularly obstructive to the host’s detriment. Aggregates of RBCs have indeed been found in the blood of most^26,27^ or a third^28^ of COVID-19 patients in three clinical studies. In a study that examined the blood of hospitalized COVID-19 patients using immunofluorescence analysis, SARS-CoV-2 spike protein punctae were found on 41% of their RBCs.^29^ *In vitro*, SARS-CoV-2 spike protein and pseudovirus attached to a nanoparticle array bearing SA derivatives.^30^ Microarray detection techniques typically fail to detect these spike protein attachments to either SA^31^ or CD147,^32^ since they are formed through nanoscale multivalent bindings.^1^

This hemagglutinating property of SARS-CoV-2 has important clinical consequences. First, with trillions of RBCs each circulating through narrow pulmonary capillaries about once per minute, even small, dynamically aggregating and disaggregating RBC clumps (as can form even in the absence of pathogens^33,34^) can impede RBC oxygenation. Peripheral ischemia, endothelial damage and vascular occlusion are indeed frequently observed in serious cases of COVID-19, as reviewed.^1,23^ In COVID-19 patients, damaged endothelium of pulmonary capillaries is often observed adjoining relatively intact alveoli,^35,36^ while hypoxemia is manifested despite normal breathing mechanics.^22,35,37-39^ These morbidities of COVID-19 parallel those of severe malaria, in which clumping of parasite-infected RBCs to other RBCs via SA terminal residues and endothelial cytoadhesion also often result in fatal outcomes.^1^

Although the blood cell types and processes entailed in clumping, clotting and vascular obstruction of COVID-19 are wide ranging, virally induced hemagglutination (HA) is a central event that is amenable to *in vitro* study. The classic HA assay was used here to study this, using cell culture supernatants and SARS-CoV-2 trimeric spike protein. Indeed, SARS-CoV-2 spike protein mixed with human whole blood caused RBC aggregation,^40^ while spike protein from two other coronavirus strains also induced HA.^41,42^ The HA assay can be applied further to study HA inhibitory effects of agents that bind to sites on spike protein, potentially shielding them from attachments to host cells. Several *in silico* studies^1^ have found that the macrocyclic lactone ivermectin (IVM) binds with high affinity to subdomains on SARS-CoV-2 spike protein, including several glycosylated binding sites.^43^ IVM achieved Nobel prize-honored distinction for success against global parasitic scourges^44^ but is of disputed efficacy in the treatment of COVID-19, as indicated, for example, by the disparity in the conclusions of this editorial^45^ and its key cited meta-analysis.^46^

The goals of this study were to determine whether principles of glycobiology as established for coronaviruses, and in particular for SARS-CoV-2, can be validated using the classic HA assay, to test whether HA inhibition is achieved by an agent indicated to competitively bind to those glycans and to determine the comparative hemagglutinating potencies of the Wuhan virus and its Alpha, Delta and Omicron B.1.1.529 variants. The clinical relevance of testing HA induced by SARS-CoV-2 using spike protein rather than whole virus and the utility of additionally testing for HA inhibition via competitive glycan binding using IVM was suggested in an earlier study.^1^ The differences in electrostatic potential of the spike protein of these four variants were studied using molecular modeling and related these to their HA-inducing potencies as experimentally observed.

## RESULTS

### Tests for Hemagglutination (HA) and for its inhibition and reversal by IVM

In the HA experiment, for the Wuhan, Alpha and Delta lineages we observed HA at a spike protein concentration of 1.06 ng/μL and above, but not below. For the Omicron spike protein, we observed HA at a minimum concentration of 0.13 ng/μL and above, but not below (Figures 1-2, Table 1).

**Table 1.**
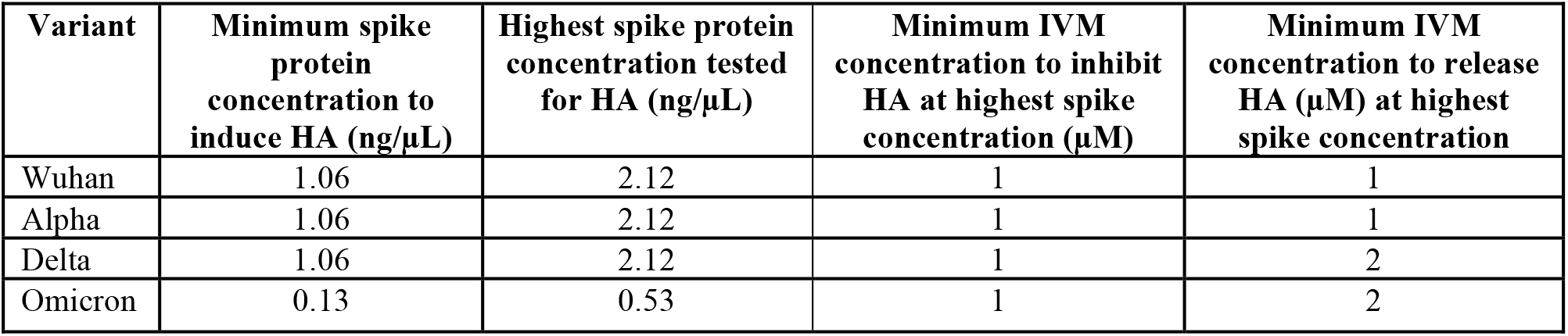
Minimum concentrations of recombinant spike protein (ng/μL) needed to induce HA when added to RBC solution, and concentrations of IVM (μM) needed to inhibit or reverse this induced HA. Tests to determine these values were each done in triplicate.

| Variant | Minimum spike protein concentration to induce HA ($\text{ng}/\mu\text{L}$) | Highest spike protein concentration tested for HA ($\text{ng}/\mu\text{L}$) | Minimum IVM concentration to inhibit HA at highest spike concentration ($\mu\text{M}$) | Minimum IVM concentration to release HA ($\mu\text{M}$) at highest spike concentration |
| --- | --- | --- | --- | --- |
| Wuhan | 1.06 | 2.12 | 1 | 1 |
| Alpha | 1.06 | 2.12 | 1 | 1 |
| Delta | 1.06 | 2.12 | 1 | 2 |
| Omicron | 0.13 | 0.53 | 1 | 2 |

**Figure 1.**
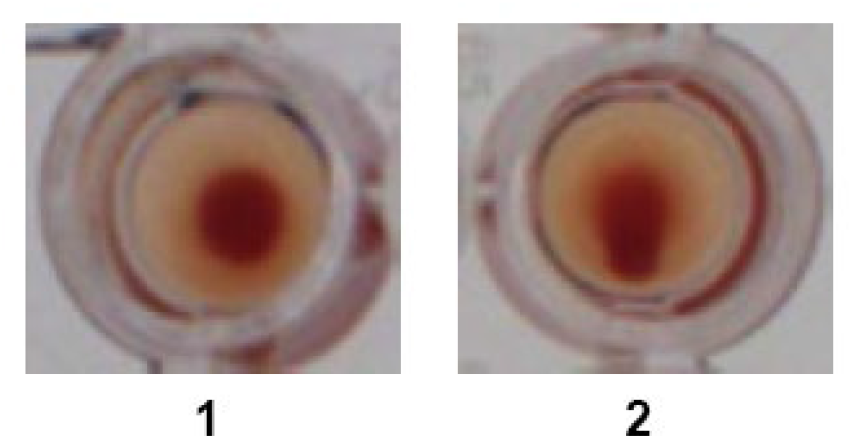
Sample images of wells in which HA occurred (1; no teardrop visible) and HA did not occur (2; teardrop visible)

**Figure 2.**
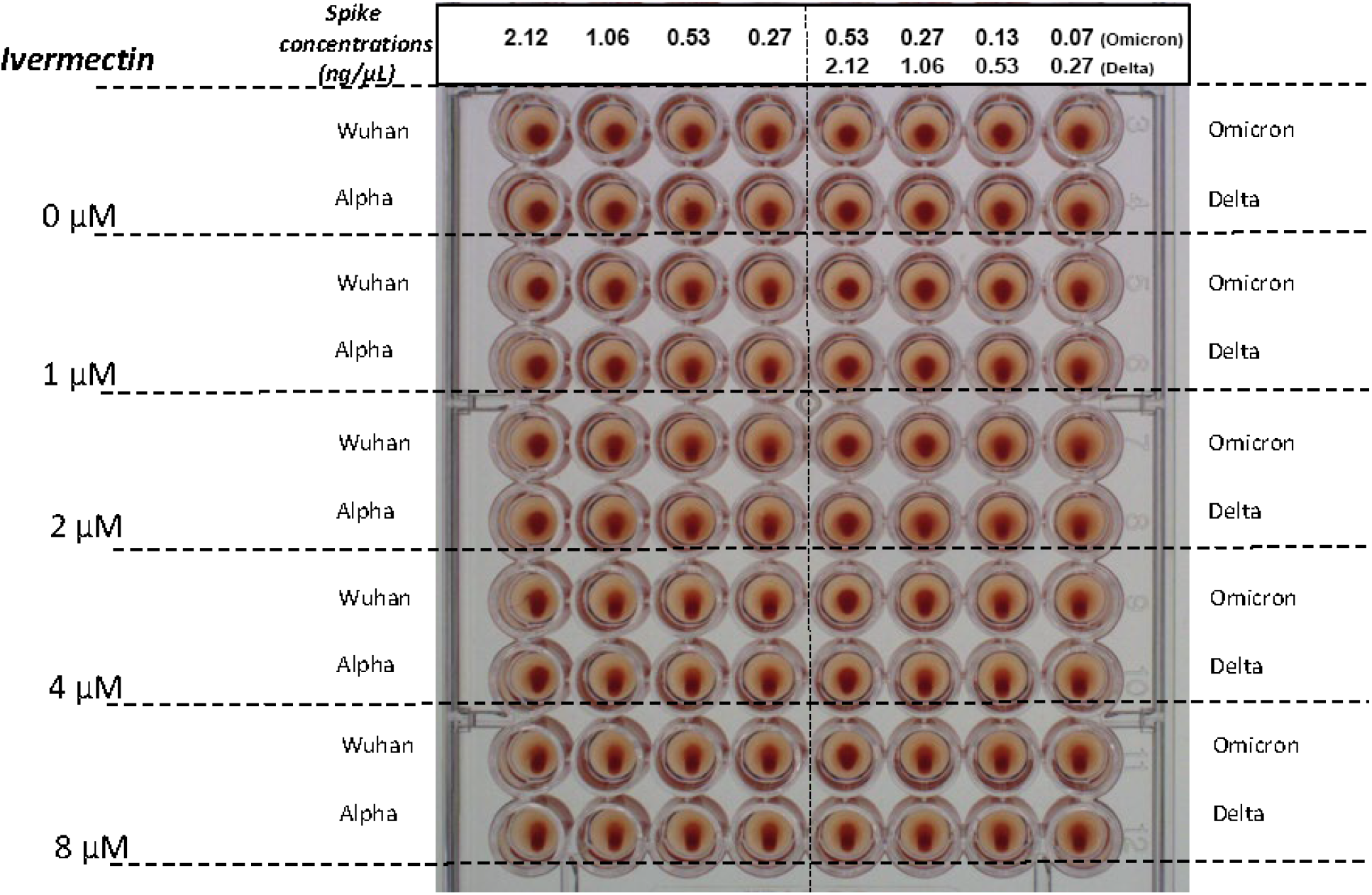
HA as induced by spike protein concentrations of 0.27, 0.53, 1.06 and 2.12 ng/μL for the Wuhan, Alpha and Delta strains of SARS-CoV-2 and at spike protein concentrations of 0.07, 0.13, 0.27 and 0.53 ng/μL for Omicron. Effects on reversal of HA are shown for IVM at concentrations of 1, 2, 4 and 8 μM added 30 min after RBCs and spike protein. Similar results were obtain for inhibition of HA by IVM (not pictured), with differences described in the Results section and summarized in Table 1.

In the HA inhibition experiment, IVM added to 2.5% RBC solution to attain a concentration of at 1 μM at 30 minutes prior to spike protein partially inhibited HA, with HA observed at a spike protein concentration of 2.12 ng/μL for the Wuhan, Alpha and Delta and of 0.27 ng/μL for the Omicron viral lineages. With IVM at 2 μM, complete inhibition of HA is observed for the Wuhan, Alpha and Delta lineages. For Omicron, IVM at 4μM is needed to totally block HA.

In the HA reversal experiment, concentrations of IVM needed to reverse HA at the highest concentration of spike tested were 1μM and 2 μM for the Wuhan/Alpha and delta/omicron viral lineages respectively.

HA was not observed with cell culture supernatants of any SARS-CoV-2 strain.

In the control experiments, RBC alone did not exhibit HA. RBCs mixed with PBS likewise did not exhibit. The addition of IVM to 50 μL of 2.5% RBCs to attain a concentration of at 8 μM did not cause hemolysis or induce HA. A solution of 2.5% DMSO and 97.5% water, the solvent for IVM, did not block or reverse HA (Figure 2 and 3).

**Figure 3.**
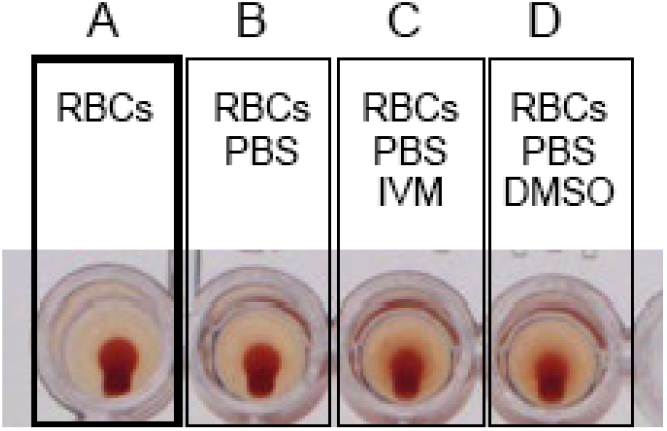
Controls: (A) RBCs alone, (B) RBCs with PBS, (C) RBCs with PBS and IVM and (D) RBCs with PBS and DMSO. No HA is observed.

### Western blot and quantification analysis

The results for quantification analysis by Western blot of spike protein in cell culture supernatants are presented in Figure 4. This shows that its spike concentration is below the concentration thresholds for induction of agglutination. Indeed for the supernatant of the Wuhan strain in cell culture at 48 h post viral infection, the concentration of N-glycosylated spike protein is approximately 0.7 ng/μL, which dilutes to half that concentration when added to wells, approximately three times lower than the minimum concentration 1.06 ng/μL of recombinant spike found to induce HA in the HA experiment described above.

**Figure 4.**
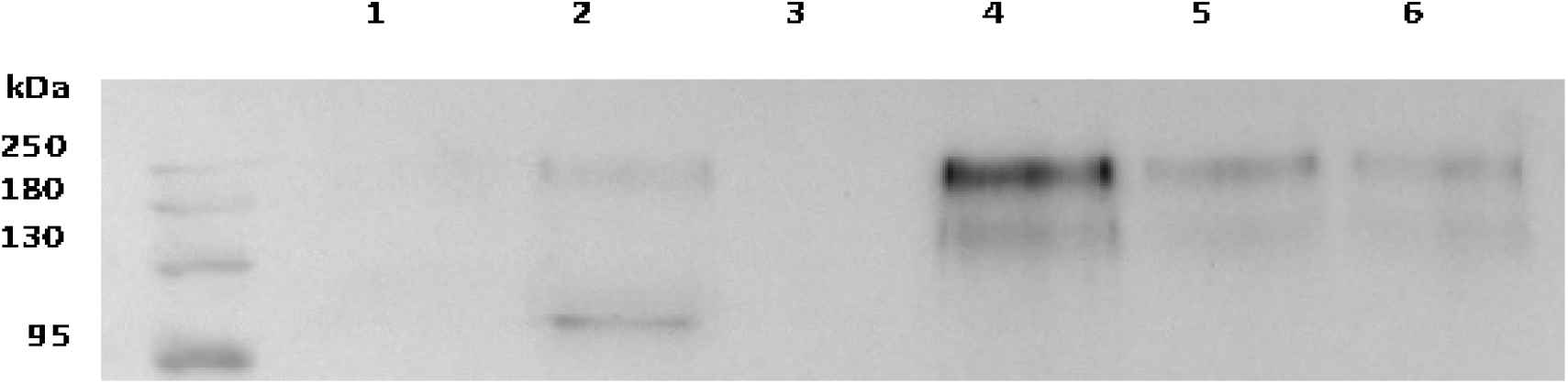
Quantification by Western blot analysis of spike protein in cell culture supernatants. Lines 1 and 2: cell culture supernatants infected with SARS-CoV-2, Wuhan strain, harvested at 24 and 48h respectively. Line 3: non-infected cell culture supernatant. Lines 4, 5 and 6: recombinant SARS-CoV-2 spike protein at 2.25, 1.25 and 0.56 ng/μL respectively.

### Molecular modeling

Since the surface of RBCs is electronegative due to a high expression of anionic sialic acids in membrane proteins and gangliosides, we used molecular modeling simulations to study the electrostatic surface potential of the spike proteins used in the present study. We focused our attention on the central area of the spike trimers which is formed by the receptor binding domain (RBD) of each monomer. The electrostatic potential of the central area of the spike trimers increased exponentially from the Wuhan initial lineage to the Omicron variant (Figure 5, lower panel). This was caused by a progressive decrease in electronegative zones (colored in red) and a concomitant increase in electropositivity (blue zones). The increase was modest between Wuhan and Alpha, larger for Delta, and reached very high levels for Omicron. As depicted in the upper panel of Figure 5, these variations in net positive electric charge between the four SARS-CoV-2 strains considered here have implications for induction of HA, since electropositivity mitigates the electrostatic repulsion between negatively charged RBC surfaces.

**Figure 5.**
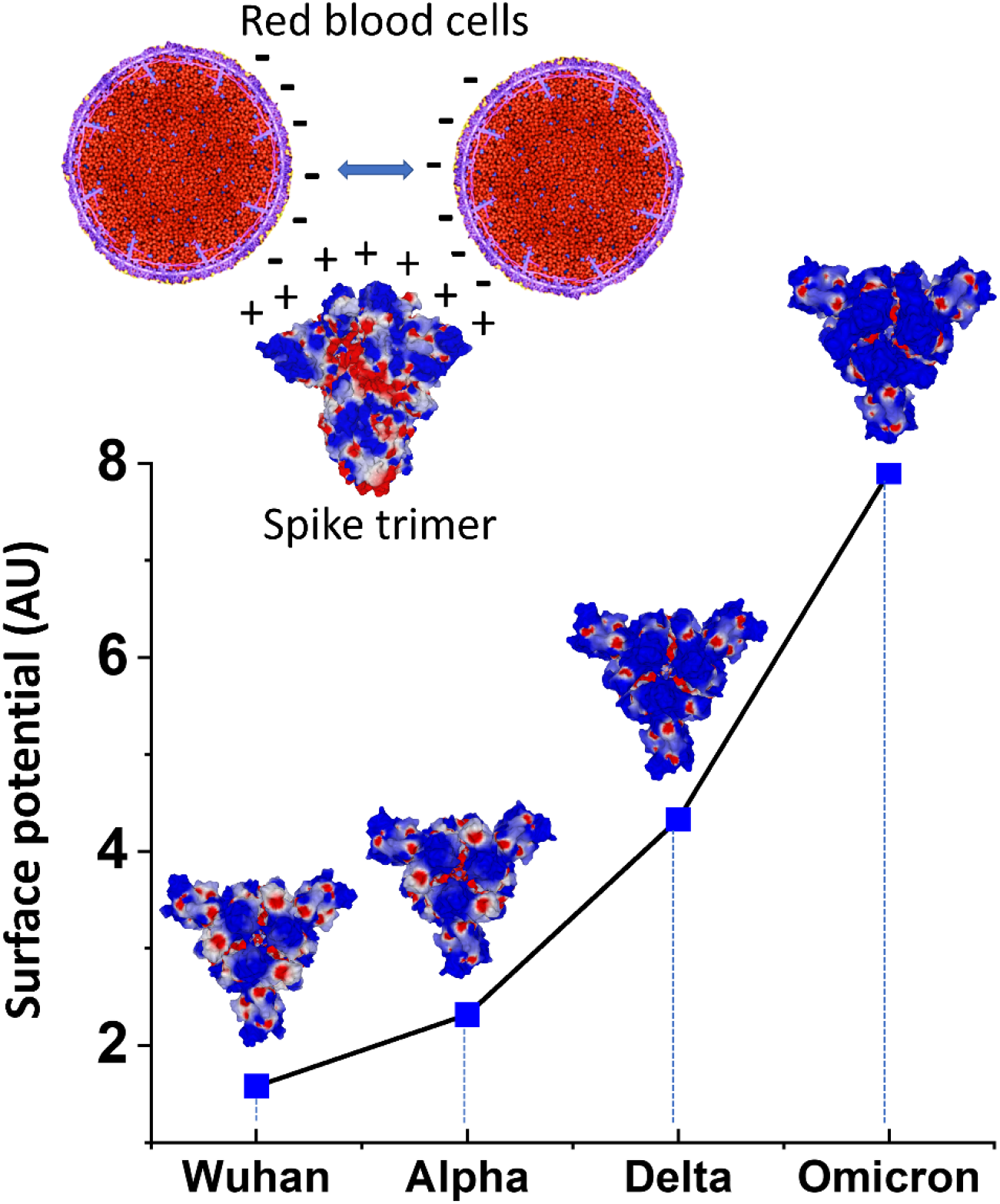
Electrostatic surface potential of SARS-CoV-2 spike trimers. Upper panel. Under physiological conditions, RBCs maintain separation from each other due to a repulsive electric zeta potential between their negatively charged surfaces. The electropositive surface of spike protein neutralizes this zeta potential, allowing closer contacts between RBCs. **Lower panel**. For all variants, the electrostatic surface potential of the spike trimer is more electropositive in the central area formed by the RBD of each monomer. Quantitative analysis of the surface potential (in AU=arbitrary units) shows an exponential increase from the Wuhan to Omicron lineages.

## DISCUSSION

For all four SARS-CoV-2 lineages tested, Wuhan, Alpha, Delta and Omicron BA.1, spike protein mixed with human RBCs induced HA. This result provides an *in vitro* counterpart to the RBC clumping found, for example, in the blood of most^26,27^ or a third^28^ of COVID-19 patients in three clinical studies, and reinforces indications that such blood cell aggregation is key to the morbidities of this disease. HA-associated vascular obstruction in COVID-19 was further demonstrated in Zebrafish embryos, which have capillary diameters^47^ and blood cell glycosylation patterns^48^ similar to those of humans. SARS-CoV-2 spike protein injected into a Zebrafish embryo vein caused the formation of small RBC clumps and an associated reduction in blood flow velocity within 3-5 minutes after injection.^49^ Also, in various *in vivo* or *in vitro* studies, SARS-CoV-2 spike protein S1 was found to cause endothelial, pulmonary and neuronal damage as well as platelet-thrombi formation and microclots.^40,50-52^

In addition to the glycan bindings reviewed above, electrostatic attraction could promote attachments between RBCs and SARS-CoV-2 spike protein underlying the observed HA. As indicated here by molecular modeling simulations, the SARS-CoV-2 spike protein central region has a net positive charge, which yields an attractive force to the negatively charged RBC surface and also mitigates the repulsive electrostatic force between RBCs. This result is consistent with prior determinations of a net positive electrostatic potential of SARS-CoV-2 spike protein.^19,53^ Regardless of whether glycan bindings or electrostatic attraction is the predominant underlying force, the HA observed here suggests, more generally, that spike protein attachments to other cells such as platelets and endothelial cells may be implicated in COVID-19 morbidities, these two cells, like RBCs, having dense surface distributions of SA-tipped glycans^1^ and corresponding negative surface charges.^16,17^ These attachments can manifest clinically in damage to endothelial cell induced by virions or free spike proteins, marked by traces of spike protein,^23,35,51^ and in the formation of fibrin-hardened clots that incorporate platelets, neutrophils and other blood cells,^54^ potentially seeded by RBC clumps.

The much stronger HA-inducing effect of spike protein of the SARS-CoV-2 Omicron variant vs. the prior lineages tested here, with thresholds of concentration for HA induction of 0.13 ng/μL for Omicron vs. 1.06 ng/μL for the Wuhan, Alpha and Delta lineages, was an intriguing result. Under the hypothesis that RBC clumping underlies the morbidities of COVID-19, especially those related to diminished efficiency of RBC oxygenation, this finding appears paradoxical, since percentages of patients having respiratory distress or requirements for oxygen or mechanical ventilation were all sharply less for Omicron as compared with prior viral lineages.^55^ Yet as noted above, increased binding affinities of viral spike protein for host cell glycoconjugates beyond optimal levels are counterproductive for infectivity, as this results in increased snagging of newly replicated virions on sialoside-binding sites of host cells and viral snagging on SA-rich non-infectious targets.^1,23,56^ The increased glycan-binding propensity of Omicron vs. prior lineages, as indicated by its increased HA-inducing activity, may thus limit the ability of an Omicron virion from an infected alveolar cell to penetrate the juncture between alveolar and capillary basal surfaces and infect an endothelial cell or otherwise enter the blood stream.

Likewise, increased net positive charge of the spike protein of Omicron vs. prior lineages could also increase viral snagging and limit penetration into the blood stream. Further confounding factors are indications from molecular modeling that while the overall electrostatic potential of Omicron spike protein is more positive than for prior lineages, its N-terminal domain (NTD) is less positive than for prior lineages,^53,57,58^ and its sialic acid binding affinity is greater.^59^ It has also been proposed, again based upon *in silico* analysis, that decreased binding affinity of mutated residues on the Omicron spike protein to a cellular receptor implicated in thromboembolic and neurological complications of COVID-19, α7nAChR, could also account for decreased morbidity of this viral variant.^60^

IVM, an electrically neutral molecule,^61^ has been found *in silico* to attach with strong affinity to 10 glycan binding sites on the spike protein of the Alpha through Delta variants,^43^ which suggests that competitive inhibition of spike protein attachments to host cell glycoconjugates could be its means of HA inhibition. IVM at concentrations of 1-2 μM for the four different viral lineages inhibited HA when added concurrently with spike protein and also reversed HA that had been induced by the prior addition of spike protein. This HA-reversal effect could account for sharp increases observed within 24 hours after administration of IVM of pre-treatment depressed SpO2 levels in severe COVID-19 patients, as summarized in Figure 6 from the most recent of three clinical studies reporting this effect.^62^ The inhibition of HA by IVM as reported here parallels the prevention of RBC clumping in Zebrafish embryos by heparan sulfate, a glycosaminoglycan likewise indicated to strongly bind to SARS-CoV-2 spike protein,^49,63^ when coinjected with spike protein.^49^ It is noteworthy that the scattered RBC clumps observed in this Zebrafish embryo study, as with those observed in the blood of COVID-19 patients,^26-28^ are much smaller than the macroscopic-scale HA—clusters of extensively interlaced RBCs—observed in this study. It is therefore likely that smaller concentrations of spike protein and IVM would be required to, respectively, induce and reverse RBC clumps of clinical relevance. Thus, the peak plasma level of IVM of approximately 412 nM as attained about four hours after a standard oral dose of 200-350 μg/kg^1^ appears to be in a range that could achieve clinical effects analogous to HA reversal observed in this study at IVM concentrations of 1-2 μM.

The HA-inducing activity of SARS-CoV-2 spike protein, which is especially potent for Omicron, raises questions as to potential risks for COVID-19 mRNA vaccines, which use spike protein as the generated antigen, even though serious adverse effects (AEs) linked to spike protein, such as myocarditis,^64-66^ are rare. Detectable levels of SARS-CoV-2 spike protein and S1 in serum or plasma have been found to persist as long as 50 days following such vaccinations.^67-69^ The possibility that spike protein migrating into the blood stream could in rare cases prompt such HA-associated AEs is suggested, for example, by a study of 1,006 subjects experiencing AEs after receiving a Pfizer/BioNTech or Moderna mRNA vaccination which found a significant degree of RBC aggregation in the blood of 948 of those subjects.^70^ These risks may be increased for younger age groups, with 301 adolescents of 13-18 years of age who received two doses of the BNT162b2 mRNA COVID-19 vaccine in one study having a 29.2% rate of cardiac AEs, ranging from tachycardia or palpitation to myopericarditis.^71^ The investigators considered chest pain, which occurred at a 4% incidence, “an alarming side effect,” however myopericarditis cases were mostly mild and temporary.

Additional experiments of interest in follow-up to this study include microscopic detection to check for initial formation of RBC clumping at spike protein concentrations lower than those which induce HA. Also, the Zebrafish study as described as above could be replicated using IVM instead of heparan sulfate as the blocking agent. Finally, HA could be tested as in this experiment but using RBCs supplemented with human serum albumin (HSA) at a physiological concentration. If IVM were to bind to spike protein glycans at the same molecular region as that which binds to HSA, that could significantly limit its HA-inhibitory effect, since 93% of IVM binds to serum proteins, mainly HSA, in blood,^72^ and that 93% of IVM would then be rendered inactive for this effect. If, on the other hand, IVM were to bind to spike protein glycans and to HSA each at different regions, then HSA, a large molecule (molecular mass of 66.5 kDa, vs. 875.1 for IVM), could considerably boost the HA-inhibitory effect of IVM through steric interference.

## MATERIALS AND METHODS

### Source and preparation of red blood cells

Red blood cells (RBCs) were provided by the “Establishment Français du Sang” (EFS) from blood bag donors qualified as “non-therapeutic blood bag” (Convention N°7828). Twenty ml of whole blood (with anticoagulant EDTA) were added to a 50 ml conical tube and filled with 30 ml of PBS at pH=7.2 before centrifugation at 800 xg for 10 min. Then, the supernatant was discarded and replaced by fresh PBS. This procedure was repeated again twice. After the last centrifugation, RBCs were diluted in PBS at a final concentration of 2.5%. RBCs were then stocked one week at 4 °C.

### Spike proteins preparation

Recombinant spike proteins (BioServUK Sheffield, United Kingdom) of these four following SARS-CoV-2 strains were used for this experiment: Wuhan, Alpha, Delta and Omicron BA.1 (BSV-COV-PR-33, BSV-COV-PR-65, BSV-COV-PR-97 and BSV-COV-OM-0.1, respectively). Four concentrations of spike protein were prepared for each of these viruses by diluting stock suspension in phosphate-buffered saline (PBS). Spike protein of each viral strain was dissolved in PBS and added to wells at final increasing concentrations of 0.27 ng/μL, 0.53 ng/μL, 1.06 ng/μL and 2.12 ng/μL for the Wuhan, Alpha and Delta lineages and of 0.07 ng/μL, 0.13 ng/μL, 0.27 ng/μL and 0.53 ng/μL for the Omicron variant.

### Cells and SARS-CoV-2 strains preparation

Vero E6 cells (ATCC-CRL-1586) were cultured as in previously described conditions^73-75^ in medium (MEM, Gibco, USA) with 2 mM L-glutamine and 10% fetal bovine serum (FBS) at 37°C in a 5% CO2 incubator. We infected Vero E6 with four viral strains genotyped by whole genome next-generation sequencing (NGS) as belonging to: Pangolin lineage B.1.1^76^ (the first major lineage following the Wuhan genotype that circulated during the first epidemic period in France, hereinafter designated “Wuhan”) and three variants: Alpha (B.1.1.7), Delta (B.1.617.2) and Omicron BA.1 (B.1.1.529). Culture supernatants were harvested 24 h post-viral infection and then passed through 0.22 μM pore-sized filters (Merck Millipore, Carrigtwohill, Ireland) to remove cell and debris and obtain viral suspension for experiments.

### Western blot and quantification analysis

Supernatants from Wuhan SARS CoV-2 infected cells prepared as described above or uninfected cells and commercial spike were lysed with 2x Laemmli Sample Buffer (#1610737, Bio-Rad, Hercules, CA, USA) with DTT (#EU0006-B, Euromedex, France) added as reducing agent and then heated at 95 °C for 5 min. Proteins were separated by 10% SDS-polyacrylamide gel electrophoresis (Laemmli, 1970) and Western-blotted on a nitrocellulose membrane. After one h of saturation in 5% nonfat dry milk with 0.3% Tween-20 in PBS, the membrane was incubated overnight with SARS/SARS-CoV-2 Coronavirus Spike Protein (subunit 1) rabbit polyclonal antibody (Thermo Fisher scientific, France) at a dilution of 1:1000 in the same buffer as for saturation. After this first incubation, the membrane was washed 10 min three times in PBS 1X-Tween buffer and then incubated for one h at room temperature with peroxydase labeled anti-rabbit donkey antibody (#NIF 824 ECL Rabbit IgG, HRP-linked whole Ab, Sigma-Aldrich Life Science, USA) diluted in saturation buffer at 1:1000. After this second incubation, the membrane was rinsed 10 min three times in PBS 1X-Tween buffer before ECL (Western Blotting Substrate, # W1001 Promega, Madison, WI, USA) revelation by image acquisition with the Fusion Fx chemiluminescence imaging system (Vilber Lourmat, Marne-la Vallée, France). Quantification in each well was calculated by measuring band intensities using ImageQuant analysis software (GE Healthcare Europe). Protein markers (New England Biolabs, #P7719S) were used for molecular mass determination.

### IVM preparation

IVM was supplied by Sigma Aldrich (St Quentin Fallavier, France). Stock solution was diluted in 2.5% of DMSO and 97.5% of water. IVM, 20 μL in volume, was added in designated wells to reach final concentrations of 1, 2, 4 and 8 μM, as specified.

### Tests for Hemagglutination (HA) and for its inhibition and reversal by IVM

Three sets of experiments were performed to test for HA induced by SARS-CoV-2 spike protein and then for HA inhibition and reversal by IVM. To test for HA, using a 96 micro-well plate, 50 μL of 2.5% RBCs in PBS was added to wells together with 62 μL of diluted spike proteins at specified concentrations. An additional 20 μL of PBS was added to attain the same total fluid volume as used in IVM inhibition and reversal experiments. This mixture was let sit for 30 min at room temperature under gentle agitation. Then, the plate was tilted for at least 30 seconds, after which, if HA had not occurred a teardrop could be observed at the bottom of the well consisting of settled RBCs (Figure 1). This teardrop was not observed if HA had occurred, i.e., if a network of linked, agglutinated RBCs had formed, as described previously.^77^ To test for inhibition of HA by IVM, 50 μL of RBCs were mixed with 20 μL of IVM at specified concentrations ranging from 1-8 μM and let sit for 30 min at room temperature under gentle agitation. Then 62 μL of spike protein was added at specified concentrations, wells were let sit for an additional 30 min at room temperature under gentle agitation, and HA was determined as in above. Finally, to test for reversal of HA by IVM, 50μL of RBCs were mixed with 62 μL of spike proteins for 30 min to determine HA as above. Then 20 μL of IVM were added at specified concentrations, wells were let sit for additional 30 min at room temperature under gentle agitation and then the plate was tilted at least 30 seconds and wells were rechecked for HA as described in above.

The following control experiments were performed. Fifty μl of RBCs deposited alone in the wells allows to check their absence of agglutination. Twenty μL of PBS were added to 50 μL of RBCs to verify absence of HA. The potential for induction of HA by IVM was tested by adding it at the highest concentration used (8 μM) to 50 μL of 2.5% RBCs. In order to test whether DMSO blocked or reversed HA, we also performed the HA inhibition and reversal experiments described respectively above using the solvent for IVM, 2.5% DMSO and 97.5% water, but without IVM.

The HA experiment was then done with viral suspensions for each viral strains by adding to the wells the same volume of viral supernatant as for commercial spike suspensions.

Each experiment was done in triplicate.

### Molecular modeling simulations

A complete structure of the reference spike protein was generated from the original 20B strain (pdb: 7bnm) as previously described.^78^ All gaps in the pdb file were fixed by inserting the missing amino acids with the protein structure prediction service Robetta [https://robetta.bakerlab.org/].^53,79^ This source file model was used to introduce the specific mutational profiles of the indicated Alpha, Delta and Omicron variants with the MUTATE tool of Swiss-PdbViewer.^53,80^ Trimeric structures in the closed pre-fusion conformation were constructed with Swiss-PdbViewer by homology with a reference model (pdb: 6VSB). All structures were then submitted to several rounds of energy minimization with the Polak-Robière algorithm.^74^ The electrostatic surface potential of the spike trimers was analyzed with Molegro and quantified with ImageJ software as described previously.^78^

## CONCLUSIONS

Spike protein from four lineages of SARS-CoV-2 induced HA in human RBCs, which supports other indications that spike protein-induced RBC clumping, as well as viral attachments to other blood cells and endothelial cells, may be key to the morbidities of COVID-19. IVM, a macrocyclic lactone indicated to bind strongly to multiple glycan sites on SARS-CoV-2 spike protein, blocked HA when added to RBCs prior to spike protein and reversed HA when added afterwards, which suggests therapeutic options for COVID-19 treatment using this drug or other competitive glycan-binding agents. The Omicron B.1.1.529 variant had significantly greater HA-inducing activity than the three prior lineages tested, which may relate in part to the finding from molecular modeling that the electrostatic charge of the central region of its spike protein was considerably more positive than for those of the prior lineages. The incongruity between Omicron’s increased HA-inducing activity and decreased morbidity may derive from limitations on virion mobility through body tissues associated with increased strength of its glycan binding or electrostatic attraction to host cell surfaces. It is not clear, however, whether this heightened HA-inducing activity of the Omicron variant might potentially increase or decrease the rate of rare HA-associated AEs for Omicron boosters vs. this rate for legacy mRNA COVID vaccines.

## Author Contributions

Conceptualization, DES, JF and BLS; methodology, JF and BLS; validation, CB, PV, JF, BLS; formal analysis, CB, BLS; investigation, AB, MM, MLB; writing— original draft preparation, DES, CB, JF, BLS; writing—review and editing, PC; supervision, JF, BLS; project administration, BLS. All authors have read and agreed to the published version of the manuscript.

## Institutional Review Board Statement

Not applicable.

## Informed Consent Statement

Not applicable.

## Conflicts of Interest

The authors declare no conflict of interest.

### Abbreviations

The following abbreviations are used in this manuscript:

ACE2: angiotensin converting enzyme 2
CD147: cluster of differentiation 147 protein, encoded by the BSG gene
COVID-19: coronavirus disease 2019
GPA: glycophorin A
NTD: N-terminal domain
RBC: red blood cell
RBD: receptor binding domain
RCT: randomized clinical trial
SA: sialic acid
SARS-CoV-2: severe acute respiratory syndrome coronavirus 2

